# Cellular and structural basis of synthesis of the unique intermediate dehydro-F_420_-0 in mycobacteria

**DOI:** 10.1101/2020.02.27.968891

**Authors:** Rhys Grinter, Blair Ney, Rajini Brammananth, Christopher K. Barlow, Paul R.F. Cordero, David L. Gillett, Thierry Izoré, Max J. Cryle, Liam K. Harold, Gregory M. Cook, George Taiaroa, Deborah A. Williamson, Andrew C. Warden, John G. Oakeshott, Matthew C. Taylor, Paul K. Crellin, Colin J. Jackson, Ralf B. Schittenhelm, Ross L. Coppel, Chris Greening

## Abstract

F_420_ is a low-potential redox cofactor used by diverse bacteria and archaea. In mycobacteria, this cofactor has multiple roles, including adaptation to redox stress, cell wall biosynthesis, and activation of the clinical antitubercular prodrugs pretomanid and delamanid. A recent biochemical study proposed a revised biosynthesis pathway for F_420_ in mycobacteria; it was suggested that phosphoenolpyruvate served as a metabolic precursor for this pathway, rather than 2-phospholactate as long proposed, but these findings were subsequently challenged. In this work, we combined metabolomic, genetic, and structural analyses to resolve these discrepancies and determine the basis of F_420_ biosynthesis in mycobacterial cells. We show that, in whole cells of *Mycobacterium smegmatis*, phosphoenolpyruvate rather than 2-phospholactate stimulates F_420_ biosynthesis. Analysis of F_420_ biosynthesis intermediates present in *M. smegmatis* cells harboring genetic deletions at each step of the biosynthetic pathway confirmed that phosphoenolpyruvate is then used to produce the novel precursor compound dehydro-F_420_-0. To determine the structural basis of dehydro-F_420_-0 production, we solved high-resolution crystal structures of the enzyme responsible (FbiA) in apo, substrate, and product bound forms. These data show the essential role of a single divalent cation in coordinating the catalytic pre-complex of this enzyme and demonstrate that dehydro-F_420_-0 synthesis occurs through a direct substrate transfer mechanism. Together, these findings resolve the biosynthetic pathway of F_420_ in mycobacteria and have significant implications for understanding the emergence of antitubercular prodrug resistance.

## Introduction

Factor 420 (F_420_) is a deazaflavin cofactor that mediates diverse redox reactions in bacteria and archaea (1). Chemically, F_420_ consists of a redox-active deazaflavin headgroup (derived from the chromophore Fo) that is conjugated to a variable-length polyglutamate tail via a phosphoester linkage (2). While the Fo headgroup of F_420_ superficially resembles flavins (e.g. FAD, FMN), three chemical substitutions in the isoalloxazine ring give it distinct chemical properties more reminiscent of nicotinamides (e.g. NADH, NADPH) (1). These include a low standard potential (−350 mV) and obligate two-electron (hydride) transfer chemistry (3, 4). The electrochemical properties of F_420_ make it ideal to reduce a wide range of otherwise recalcitrant organic compounds (5–7). Diverse prokaryotes are known to synthesize F_420_, but the compound is best characterised for its roles in methanogenesis in archaea, antibiotic biosynthesis in streptomycetes, and metabolic adaptation of mycobacteria (1, 8–11). In mycobacteria, F_420_ is involved in a plethora of processes: central carbon metabolism, cell wall synthesis, recovery from dormancy, resistance to oxidative stress, and inactivation of certain bactericidal agents (7, 12–14). In the human pathogen *Mycobacterium tuberculosis*, F_420_ is also critical for the reductive activation of the newly approved clinical antitubercular prodrugs pretomanid and delamanid (15–17).

Following the elucidation of the chemical structure of F_420_ in the 1970s, the F_420_ biosynthesis pathway in archaea was determined through a combination of in situ biochemistry and recombinant protein analysis (1, 2). Described briefly, the deazaflavin fluorophore Fo is synthesized through condensation of 5-amino-6-ribitylamino-2,4(1H,3H)-pyrimidinedione and L-tyrosine by the SAM-radical enzymes CofG and CofH (18). The putative enzyme CofB synthesizes 2-phospholactate (2PL), which links Fo to the glutamate tail of mature F_420_ (19). Subsequently, the nucleotide transferase CofC condenses 2PL with GTP to form the reactive intermediate L-lactyl-2-diphospho-5’-guanosine (LPPG) (20). The phosphotransferase CofD then transfers 2PL from LPPG to Fo, leading to the formation of F_420_-0 (i.e. F_420_ with no glutamate tail) (21). Finally, the GTP-dependent glutamate ligase CofE adds a variable-length γ-linked glutamate tail to produce mature F_420_ (22, 23). With the exception of the putative lactate kinase CofB, the enzymes responsible for F_420_ biosynthesis in archaea have been identified and characterized to varying extents (1). Crystal structures have been obtained for CofC, CofD and CofE from methanogenic archaea, providing some insight into how these enzymes function, but questions surrounding their catalytic mechanisms remain unresolved (23–25). For example, the crystal structure of CofD from *Methanosarcina mazei* was solved in the presence of Fo and GDP; however, no divalent cation(s) required for catalysis were present in the structure and the ribosyl tail group of Fo, which receives the 2PL moiety from LPPG was disordered, precluding an understanding of the catalytic mechanism of this step in F_420_ biosynthesis (21, 25).

It was assumed that the biosynthesis pathway for archaeal F_420_ was generic to all F_420_ producing organisms (1). However, recent studies have shown that the structure and biosynthesis of F_420_ varies between producing organisms (24, 26). F_420_ produced by the proteobacterial fungal symbiont *Paraburkholderia rhizoxinica* was found to incorporate 3-phospho-D-glycerate (3PG) in the place of 2PL, producing a chemically distinct F_420_ (26). In parallel, analysis of purified F_420_ biosynthesis enzymes from mycobacteria indicated that the central glycolytic and gluconeogenic intermediate phosphoenolpyruvate (PEP), rather than 2PL, is a precursor for F_420_ biosynthesis (24). In contrast to *P. rhizoxinica*, in mycobacteria, mature F_420_ is chemically analogous to that produced by archaea (27). All mycobacterial species possess the four enzymes required for F_420_ biosynthesis. However, as these enzymes catalyze reactions distinct to their archaeal homologues, the following alternative nomenclature is applied compared to the archaeal enzymes: FbiD (homologous to CofC), FbiC (a single protein with domains homologous to CofG and CofH), FbiA (homologous to CofD) and FbiB (N-terminal domain homologous to CofE) (8). In addition to its CofE-like domain, FbiB possesses an FMN-binding C-terminal domain and biochemical evidence suggests it is responsible for the reduction of the moiety derived from PEP (24, 28).

However, several findings have cast doubt on whether the proposed revised biosynthesis pathway of F_420_ is physiologically relevant. The predicted use of PEP in F_420_ biosynthesis in mycobacteria would lead to the formation of the oxidized intermediate compound dehydro-F_420_-0 (DH-F_420_-0). The production of DH-F_420_-0 was detected in a coupled enzyme assay containing purified FbiD and FbiA with PEP supplied as the substrate but has yet to be detected in mycobacterial cells (24). The study also showed that CofC from *Methanocaldococcus jannaschii* utilized PEP rather than 2PL for F_420_ biosynthesis (24), leading the authors to conclude that PEP is the general precursor for F_420_ biosynthesis in prokaryotes. However, these findings contradict previous analysis of CofC activity in *M. jannaschii* cell lysates (19, 21), as well as recent biochemical analysis, which shows that CofC preferentially utilizes 2PL for F_420_ biosynthesis (26). In turn, these findings cast doubt on whether PEP is truly the preferred substrate for mycobacterial F_420_ biosynthesis and whether DH-F_420_-0 is the physiological intermediate in this pathway.

In this work, we first resolved this ambiguity by analyzing the F_420_ biosynthetic pathway in *M. smegmatis* in whole cells. We demonstrate that PEP, not 2PL, is the substrate for F_420_ biosynthesis in mycobacterial cells, suggesting that divergent biosynthesis pathways are utilized to generate F_420_ in different prokaryotic species. Consistent with this result, we determine that DH-F_420_-0 is the physiological intermediate for F_420_ biosynthesis in mycobacteria, and is present in high quantities in cells lacking FbiB and comes bound to FbiA purified from *M. smegmatis*. Furthermore, to elucidate the catalytic mechanism for the formation of the novel intermediate DH-F_420_-0, we determined the crystal structure of FbiA in the presence and absence of its substrate and product compounds. These data resolve long-standing questions about the catalytic mechanism of FbiA and CofD in F_420_ biosynthesis. Moreover, they provide a target for therapeutic intervention through the inhibition of F_420_ biosynthesis, as well as insight into potential mechanisms for the emergence of delamanid and pretomanid drug resistance through mutations in FbiA.

## Results

### Phosphoenolpyruvate is the substrate for the biosynthesis of F_420_ in mycobacterial cells

To determine whether PEP or 2PL is the substrate for F_420_ biosynthesis in mycobacteria (Figure 1A), we spiked clarified cell lysates from *M. smegmatis* with GTP and either PEP or 2PL, and monitored the synthesis of new F_420_ species through HPLC coupled with fluorescence detection. In cell lysates spiked with PEP, a species corresponding to F_420_-0 in the F_420_ standard was present, which was absent from both the untreated and 2PL spiked lysates (Figure 1B). The formation of this F_420_-0-like species in PEP spiked lysates corresponded to a decrease in Fo levels, suggesting that synthesis of DH-F_420_-0 from PEP is occurring (Figure 1B). These data strongly suggest that PEP, not 2PL, is the precursor for F_420_ biosynthesis in *M. smegmatis*.

**Figure 1.**
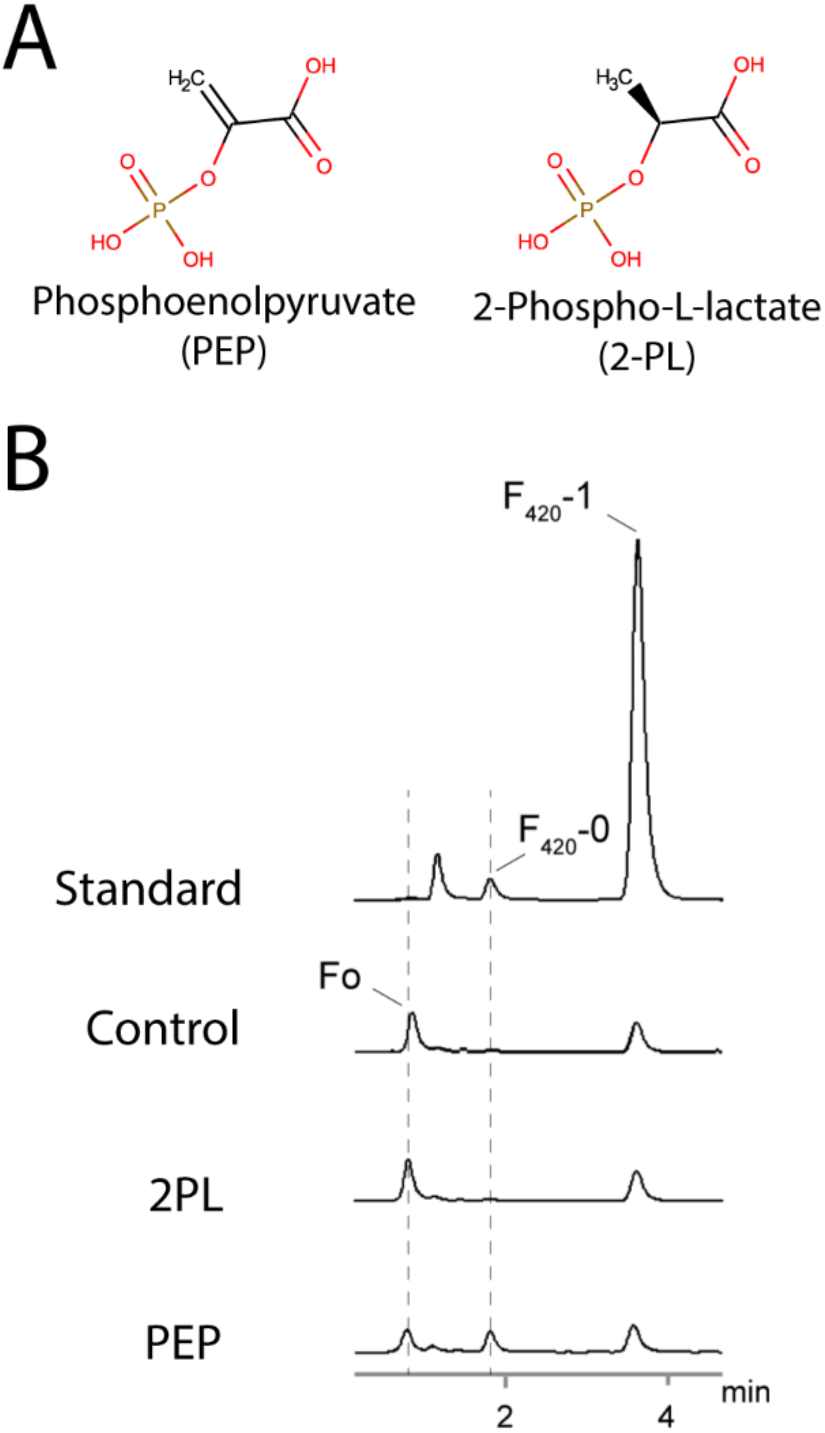
PEP but not 2PL stimulates DH-F_420_-0 synthesis in *M. smegmatis* cell lysates. (A) 2D structures of PEP and 2PL demonstrating the difference (double bond or single bond) in bonding between carbon 2 and 3. (B) Fluorescence emission detection chromatogram from HPLC of *M. smegmatis* lysates spiked with either 2PL, PEP or an unspiked control. Synthesis of a species with characteristic F_420_ fluorescence (Ex: 420 nm, Em:480 nm) corresponding to F_420_-0 from the purified standard was only detected in the PEP-spiked lysate. The appearance of this F_420_-0-like species coincided with a decrease in the presence of Fo, suggesting that PEP is the precursor for F_420_ synthesis in *M. smegmatis* in cells. F_420_-1 in the standard corresponds to F_420_ with a single glutamate moiety.

While the lysate spiking experiment establishes that PEP is specifically utilised for F_420_ synthesis in *M. smegmatis*, the fluorescent detection method utilised does not chemically differentiate between F_420_-0 or DH-F_420_-0. As PEP is utilised, it would be expected that DH-F_420_-0 is produced. However, DH-F_420_-0 may be rapidly reduced to F_420_-0 rather than accumulating in the cell. To confirm the synthesis of the DH-F_420_-0 in *M. smegmatis*, we created isogenic deletions in the four F_420_ biosynthesis genes: *fbiD*, *fbiC*, *fbiA*, and *fbiB* (Figure 2A). The genome sequences of these deletion strains were determined, confirming clean deletion with no secondary mutations present. We then detected the deazaflavin species present in clarified cell lysates from these strains using fluorescence-coupled HPLC and LC-MS. As expected, based on the proposed function of these enzymes, mature F_420_ was only detected in wild-type cell lysates and possessed a polyglutamate tail length of three to eight (Figure 2B, Figure S1). Fo was detected in the wildtype and all mutants with the exception of *ΔfbiC*, consistent with the function of this enzyme in the synthesis of the Fo-deazaflavin moiety (Figure 2A, C, Figure S1). The proposed biosynthetic intermediate DH-F_420_-0 was detected only in cell lysates of the *ΔfbiB* strain (Figure 2D, Figure S1). No F_420_-0 was detected in wildtype or mutant strains.

**Figure 2.**
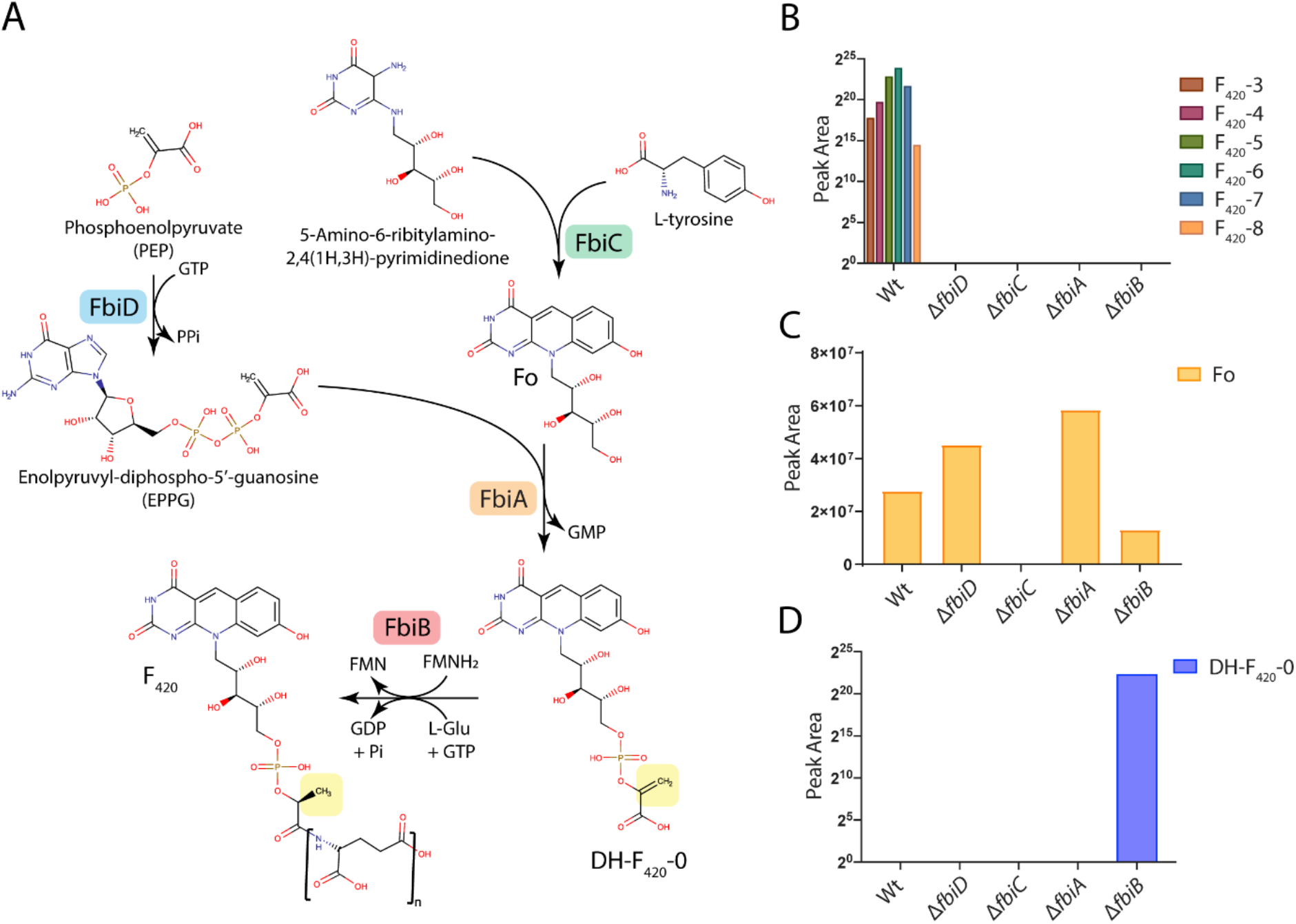
Mutagenic dissection of the F_420_ biosynthesis pathway in *M. smegmatis* reveals DH-F_420_-0 is the biosynthetic intermediate in mycobacteria. (A) A schematic of the F_420_ biosynthesis pathway in *M. smegmatis* with PEP, rather than 2PL, utilised by FbiD to create the reaction intermediate EPPG. The enzymes responsible for catalytic steps are shown, along with the 2D-structures of proposed pathway intermediates and mature F_420_. The yellow box highlights the reduction of DH-F_420_-0, proposed to be mediated by the C-terminal domain of FbiB using FMNH2. LC-MS detection of mature F_420_ species (B), Fo (C) and DH-F_420_-0 (D) in *M. smegmatis* cell lysates of wildtype and F_420_ biosynthesis pathway mutants confirming the proposed function of the F_420_ biosynthetic genes detecting the novel intermediate DH-F_420_-0 in whole cells. F_420_-X species in panel B correspond to different lengths of the ploy-glutamate chain where X = n tail length.

The presence of DH-F_420_-0 (and absence of detectable F_420_-0) in whole cells demonstrates that it is the central physiological intermediate in mycobacterial F_420_ biosynthesis. This also lends support to the biochemical and cellular assays indicating that PEP, not 2PL, is the substrate for this pathway in mycobacteria. Furthermore, in addition to its role as the F_420_ glutamyl-ligase, structural and biochemical analysis suggests that FbiB is responsible for the reduction of DH-F_420_-0 (24, 28). The detection of DH-F_420_-0 only in the *ΔfbiB* strain demonstrates that this intermediate is rapidly turned over in the cell and supports the hypothesis that FbiB and not another enzyme performs this step in mycobacterial F_420_ biosynthesis.

### FbiA co-purifies with its product dehydro-F_420_-0

In order to determine the catalytic mechanism for the synthesis of the novel intermediate DH-F_420_-0, we overexpressed and purified FbiA from *M. smegmatis*. Purified FbiA from *M. smegmatis* possessed a light-yellow color, indicating co-purification with a product or substrate molecule (Figure S2A). The nature of this substrate was investigated using fluorescence spectroscopy, with purified FbiA found to have a broad absorbance peak at 400 nm and a corresponding emission peak at 470 nm (Figure S2B), which is consistent with the presence of a deazaflavin with a protonated 8-OH group (16). We then utilized LC/MS to identify the deazaflavin species associated with FbiA and found the major species was its product DH-F_420_-0 (Figure S2C). In addition, significant quantities of mature F_420_ species were also associated with FbiA, suggesting that it also binds to mature F_420_ present in the cytoplasm (Figure S2C).

### The crystal structure of FbiA reveals an active site with open and closed states

In order to resolve the catalytic mechanism of DH-F_420_-0 synthesis, purified FbiA was crystallized and its structure was determined at 2.3 Å by X-ray crystallography (Table S1). FbiA crystallized as a dimer mediated by the interaction of three α-helices and a β-sheet (Figure 3A). This dimer is predicted to be stable by the protein-interaction prediction program PISA (29) and the molecular weight of FbiA determined by SEC-MALS shows it forms a dimer in solution (Table S2, Figure S2D). The structure of CofD from *M. mazei*, a homologous enzyme that instead utilizes LPPG derived from 2PL as its substrate, also crystallised as a dimer with an analogous interface to FbiA (25). Despite the copurification of FbiA with DH-F_420_-0, only weak electron density attributable to DH-F_420_-0 was observed in the catalytic site of molecule B (Mol. B) of the FbiA dimer (Figure S3). To obtain the product bound structure of FbiA, DH-F_420_-0 was purified from recombinant FbiA and soaked into existing crystals of FbiA. Using this procedure, electron density clearly attributable to the Fo and phosphate moieties of DH-F_420_-0 was observed in Mol. B of FbiA, allowing modelling of the product bound structure (Figure S3). Density for the carbonyl group of DH-F_420_-0 was less well resolved, suggesting it exists in multiple conformations in product-bound FbiA (Figure S3). Similarly, FbiA crystals were soaked with Fo and GDP, individually or in combination, and structures of substrate-bound FbiA were determined (Figure S3, Table S1).

**Figure 3.**
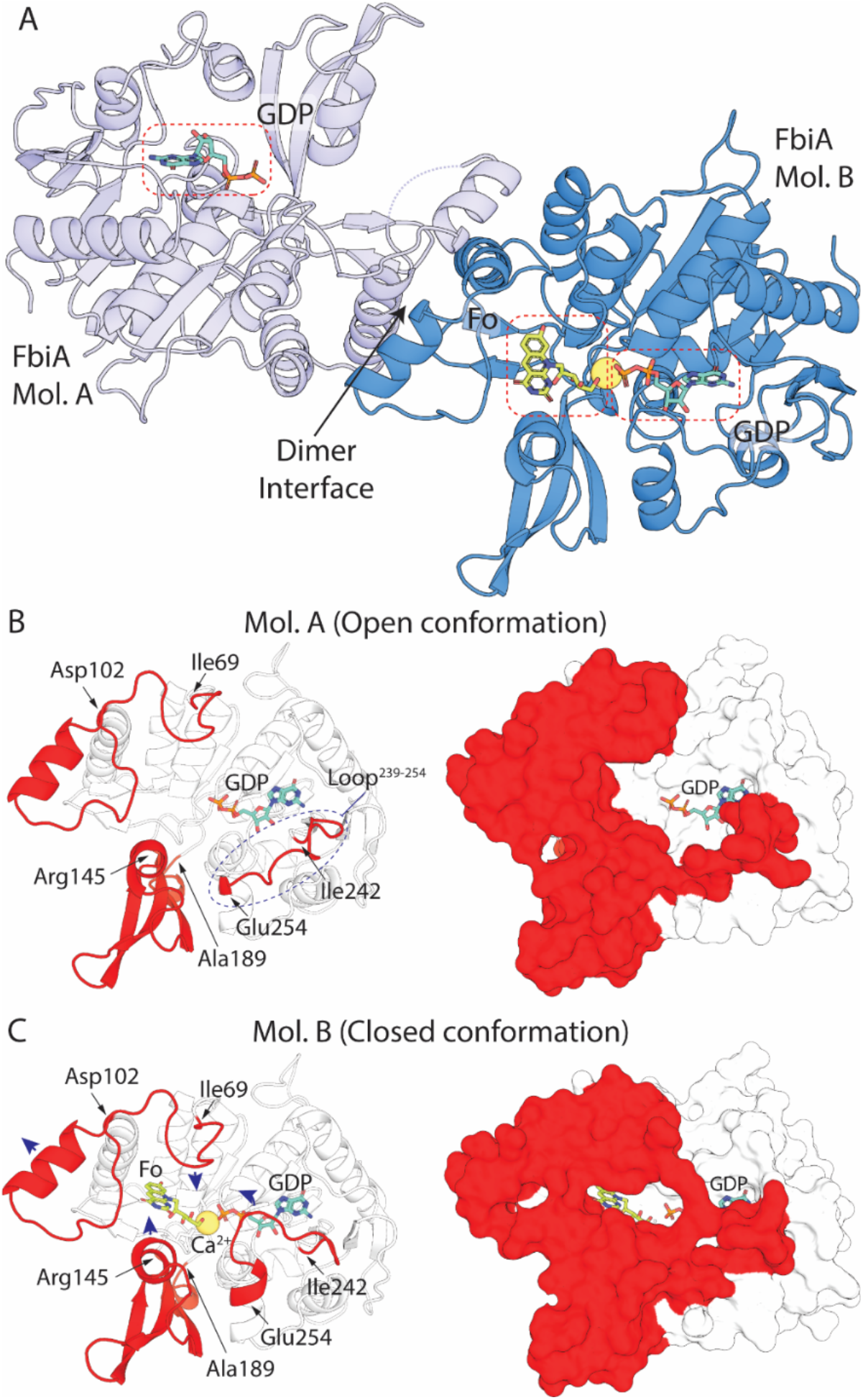
The crystal structure of FbiA captures the enzyme in open and closed states. (A) The crystal structure of FbiA from *M. smegmatis* in complex with Fo and GDP. FbiA is shown as a cartoon representation with Mol. B colored in sky blue and Mol. A colored in light blue. GDP and Fo are shown as a stick representation and Ca^2+^ is shown as a yellow sphere. (B) Mol. A from the FbiA structure exists in an open conformation. Left shows Mol. A as a cartoon with loops and subdomains which differ in conformation in Mol. B highlighted in red. Right shows Mol. A as a surface representation with mobile regions highlighted in red. (C) Mol. B. of FbiA structure exists in a closed ‘catalytically ready’ state. Left is displayed as panel B, with the direction of movement of loops compared to Mol. A shown with blue arrows. Right shows Mol. B as in panel B, demonstrating how the mobile regions enclose the FbiA active site.

Comparison of the Mol. A and B from the FbiA dimer reveals the active site of the enzyme in distinct open and closed conformations (Figure 3). In Mol. A, the active site is locked in an open state due to participation of an extended loop (amino acids 239-254) in crystal packing (Figure 3B). In this open state, FbiA has a lower apparent substrate affinity, with no density attributable to Fo or DH-F_420_-0 and only weak density for GDP observed in the respective co-crystal structures (Figure S3). In contrast, in Mol. B the extended loop (AA 239-254) is partially disordered in the non-GDP bound structures and encloses GDP in the active site in GDP-bound structures (Figure 3C). In Mol. B, additional conformational changes are observed in amino acids 69-102 and a subdomain composed of amino acids 145-189, creating the binding pocket for the deazaflavin moiety of Fo or DH-F_420_-0, which is not present in Mol. A (Figure 3B, C). The conformational differences observed between Mol. A and Mol. B are consistent in the apo, substrate and product-bound structures, demonstrating they are not substrate-induced and are likely representative of conformational differences of the enzyme in solution.

### The crystal structures of FbiA in substrate- and product-bound forms provide mechanistic insight into dehydro-F_420_-0 synthesis

The resolution of the structure of FbiA in the presence of its substrate and product compounds provides key insights into the catalytic mechanism of this unique phosphotransferase. It was previously established that FbiA and its archaeal homologue CofD require the presence of the divalent cation Mg^2+^ for activity (21, 24). However, it remained to be resolved whether Mg^2+^ is bound stably in the FbiA active site during catalysis and the mode of coordination of the ion(s). In the GDP-bound structures of FbiA, a single metal ion was present in the active site of Mol. B of FbiA (Figure 4A, Figure S4A, Figure S5). Interestingly, no metal ion was observed in Mol. A, despite the presence of GDP, suggesting that its recruitment is conformation-dependent (Figure S2). As calcium acetate is present at high concentration (0.2 M) in the crystallization condition, and magnesium is absent, we modelled this ion as Ca^2+^. While the activity of FbiA in the presence of Ca^2+^ has not been tested, it has been shown to be interchangeable with Mg^2+^ in some phosphotransferase and phosphohydrolase reactions (30, 31). In the GDP-only structure, the Ca^2+^ ion is directly coordinated by aspartates 45 and 57, an oxygen atom of the β-phosphate of GDP, two H_2_O molecules, and a glycerol molecule (Figure S4B). Aspartate 57 exhibits bidentate coordination of the Ca^2+^ ion leading to a coordination number of seven with distorted octahedral geometry.

**Figure 4.**
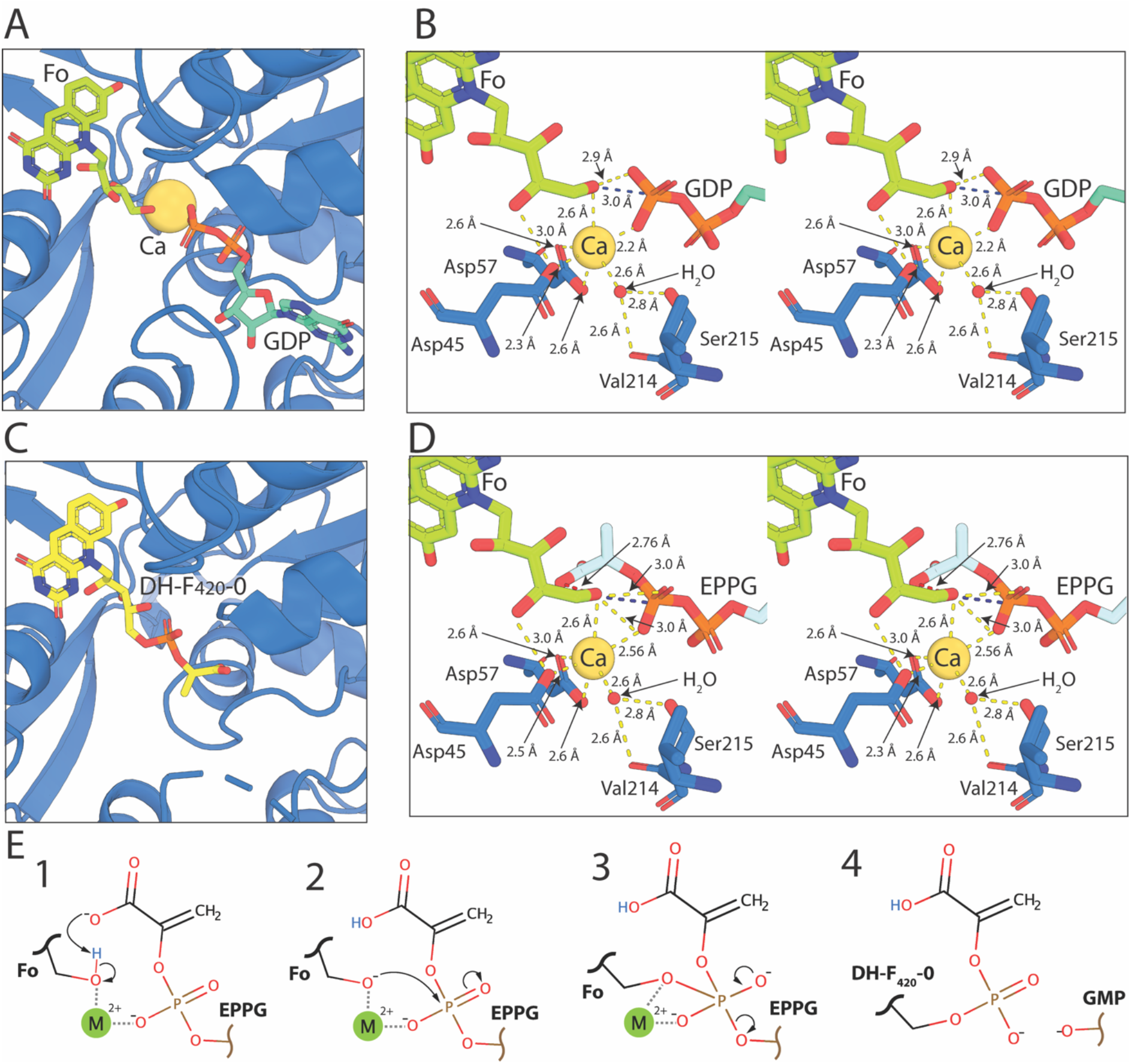
Resolution of the structure of FbiA in the presence of Fo, GDP and DH-F_420_-0 provides insight into its catalytic mechanism. (A) Fo and GDP in complex with Mol. B of FbiA in coordination with the catalytic Ca^2+^ ion. FbiA is shown as a sky-blue cartoon, Fo and GDP as sticks and Ca^2+^ as a sphere. (B) A stereoview of the catalytic center of the FbiA active site in complex with Fo and GDP, showing FbiA sidechains involved in coordinating the catalytic metal ion and a coordinating H_2_O molecule. Bond distances <3.2 Å are shown as yellow dashed lines, the distance between the terminal OH of Fo and P of the β-phosphate of GDP is highlighted in blue. (C) DH-F_420_-0 in complex with FbiA, shown as in panel A. (D) A stereoview of the FbiA catalytic centre with the reaction substrate EPPG model in place of GDP displayed in panel C, with the close proximity between the carboxylic acid group of EPPG and the terminal OH of Fo highlighted with a red dashed line. (E) A schematic showing the proposed catalytic mechanism for the formation of DH-F_420_-0 by FbiA.

In the GDP and Fo bound structure, the coordination of Ca^2+^ is analogous to the GDP-only structure. However, the glycerol molecule and one of the H_2_O molecules observed in the GDP only structure are displaced by the ribosyl chain of Fo, resulting in a coordination number of six with octahedral geometry (Figure 4B, Figure S4B). The terminal hydroxyl group of Fo is significantly closer to the Ca^2+^ ion (2.6 Å) and to the β-phosphate of GDP (2.8 Å from O, and 3.0 Å from P) than the coordinating hydroxyl of glycerol, which is not within bonding distance of GDP. These bond distances between the hydroxyl of Fo and GDP, as well as the central orientation of the hydroxyl of Fo towards the β-phosphate of GDP, place it in an ideal position to act as the acceptor substrate for the transfer of PEP catalysed by FbiA (Figure 4B). In the Fo and DH-F_420_-0 bound structures, no density corresponding to a Ca2+ ion was observed (Figure 4C, Figure S4C). This suggests that binding of FbiA to its catalytic metal ion is contingent on complex formation with enolpyruvyl-diphospho-5’-guanosine (EPPG) (Figure 2A), which is substituted for GDP in our structures due to the instability of the F_420_ pathway intermediate (24). The ability of FbiA to bind Fo in the absence of GDP and Ca^2+^ suggests that substrate binding to FbiA is non-sequential; however, recruitment of all three components is required for catalysis to proceed.

Based on these structural data and previous biochemical characterization of FbiA and CofD we propose a catalytic mechanism for synthesis of the DH-F_420_-0 (Figure 4E) (21, 25). Our structural data agree with previous work that suggests that CofD does not form a covalent intermediate as part of the reaction mechanism (21), but rather the reaction proceeds through direct nucleophilic attack of the β-phosphate of EPPG by the terminal hydroxyl of Fo. This leads to the formation of a pentavalent transition state between Fo and EPPG that is stabilized by the catalytic metal ion (Figure 4E). In order for the hydroxyl group of Fo to perform nucleophilic attack, it needs to be activated through deprotonation. The carboxylic acid group of EPPG is a likely candidate for this activation, as it is the only acidic group in close proximity to the hydroxyl group of Fo when EPPG is modelled in the FbiA structure in place of GDP (Figure 4D). Additionally, the activation of Fo by the carboxylic acid group of EPPG would provide FbiA with substrate specificity for EPPG over GDP and GTP. Following the formation of the pentavalent reaction intermediate, GMP would act as the leaving group, leading to the formation of the phosphodiester bond between PEP and Fo and the formation of DH-F_420_-0 (Figure 4E).

## Discussion

Integrating these findings with other recent literature, it is now clear that the substrate for the initial stage of F_420_ tail biosynthesis differs between F_420_ producing organisms (19, 24, 26). We definitively show here that, in mycobacteria, PEP is the substrate for F_420_ biosynthesis, resolving the ambiguity in the literature (24, 26). In contrast, in the archaeal and proteobacterial species that have been analysed, 2PL and 3PG respectively are preferentially utilized (19, 26). This divergent substrate utilization occurs despite the enzymes responsible for this stage of synthesis (FbiD/CofC and FbiA/CofD) sharing a common evolutionary history (8). This suggests that the substrate specificity of these enzymes has evolved in response to selection to maintain compatibility between the substrate used for F_420_ biosynthesis and what is available in the cellular metabolite pool. Both PEP and 3PG are intermediates in central metabolic pathways, including glycolysis, whereas 2PL is not thought to be present in significant quantities in most organisms (24, 32, 33). This makes PEP and 3PG compatible substrates for F_420_ biosynthesis in *Mycobacterium* spp. and *P. rhizoxinica* respectively, with the specifics of cellular metabolism of each organism likely dictating which compound was selected for F_420_ biosynthesis. In contrast, in the archaeon *Methanobacterium thermoautotrophicum,* 2PL is present at micromolar concentrations (19). However, in archaeal species, it remains to be determined how 2PL is synthesized and whether this compound plays a wider role as a general metabolite beyond F_420_ biosynthesis.

Phylogenetic analysis of FbiD/CofC and FbiA/CofD suggests that these proteins were horizontally transferred from bacteria and archaea (8). Based on this analysis, it is curious that mycobacteria reduce DH-F_420_-0 produced via PEP to F_420_, rendering it chemically identical to that produced with 2PL. The redox properties of the deazaflavin group of DH-F_420_ and F_420_ are identical and chemically the molecules are very similar, posing the question: why is reduction of DH-F_420_-0 is required? A plausible explanation is that actinobacteria originally utilized 2PL for F_420_ synthesis, with a switch to PEP occurring at a later stage in evolution. As a result, the F_420_-dependent enzymes present in mycobacteria evolved to recognize the non-planar 2PL moiety of F_420_, requiring reduction of DH-F_420_ to maintain compatibility after the substrate switch. Previous structural and biochemical analysis suggests the C-terminal domain of mycobacterial FbiB is responsible for the reduction of DH-F_420_-0 (24, 28). This domain is present in all mycobacterial species, but is absent from FbiB/CofE in most other F_420_ producing organisms, including *M. mazei* and *P. rhizoxinica*, that produce F_420_ through pathways that do not require this reductive step (8, 26). This conclusion is supported by our cellular analysis of F_420_ biosynthesis in *M. smegmatis*, which shows that DH-F_420_-0 accumulates in the *ΔfbiB* strain (Figure 2A, D).

The structural analysis of FbiA that we present in this work provides unprecedented insight into the catalytic mechanism for the novel phosphotransferase reaction employed at this step in F_420_ biosynthesis. The crystal structure of FbiA shows that this enzyme employs a flexible active site to capture Fo and EPPG, precisely positioning them for catalysis. Determination of the fully resolved FbiA substrate complex, in the presence of a single catalytic metal ion, provides a clear picture of the mechanism of catalysis of this enzyme. In this structure, the terminal hydroxyl group of Fo is ideally positioned for nucleophilic attack of the β-phosphate of EPPG, strongly suggesting that DH-F_420_-0 biosynthesis occurs through direct transfer of PEP to Fo, via a pentavalent phosphate intermediate that is stabilized by the catalytic metal ion. The positioning of the Fo terminal hydroxyl group in our structure in relation to EPPG is strikingly similar to that of the attacking ribose in the final step in DNA ligation by T4 ligase, recently resolved by X-ray crystallography (34). This is consistent with both reactions resulting in the formation of a phosphodiester bond through direct nucleophilic attack of a diphosphonucleoside intermediate. As no acidic side chains are present in proximity the terminal hydroxyl of Fo in our structure of FbiA, it is likely that deprotonation of this group for nucleophilic attack is EPPG induced, possibly by the PEP carboxyl moiety. These data are also consistent with biochemical analysis of CofD from *M. mazei*, which did not detect the formation of a catalytic reaction intermediate during the synthesis of F_420_-0 (21).

The resolution of the F_420_ biosynthesis pathway also has implications for tuberculosis treatment. It has been proposed that F_420_ biosynthesis represents a promising target for the development of drugs for the treatment of *M. tuberculosis* given the pleiotropic role of this cofactor and its absence from human cells. Given FbiA mediates the key step in F_420_ biosynthesis, the structural insights into FbiA catalysis provide a basis for the development of inhibitory compounds targeting F_420_ biosynthesis. In addition, loss-of-function of FbiA causes resistance to the clinical nitroimidazole prodrugs delamanid and pretomanid, both of which are activated by the F_420_H_2_-dependent reductase Ddn (15, 17, 35). Hence, the structural and mechanistic insights provided here will enable prediction of which substitutions are likely to impair or inactivate FbiA, thus conferring resistance to these compounds.

## Methods

### Creation of *M. smegmatis* F_420_ biosynthesis mutant strains

*M. smegmatis* MSMEG_5126 was deleted in wild-type *M. smegmatis* mc^2^ 155 using a two-step allelic replacement strategy. Two 0.8 kb fragments containing sequences from the left and right flanking regions of the MSMEG_5126 (*fbiC*) gene were cloned as separate constructs and later combined to make the deletion construct. The left flanking fragments were amplified using ProofStart DNA polymerase (Qiagen) with primers MSMEG_5126left and MSMEG_5126leftrev, and the PCR product was subsequently cloned into the SacI/BamHI sites of pUC18, creating plasmid pUC-MSMEG_5126left (primers described in Table S3). A 0.8 kb fragment containing sequence from the right side of MSMEG_5126 was amplified using primers MSMEG_5126right and MSMEG_5126rightrev and cloned into the XbaI/BamHI sites of pUC18, creating plasmid pUC-MSMEG_5126right. The right flanking sequence was then excised from pUC-MSMEG_5126right using XbaI/SacI and subcloned into Xba/SacI-digested pUC-MSMEG_5126left, fusing the left and right flanking sequences to create plasmid pUC-ΔMSMEG_5126.

The 1.6 kb fused insert was then liberated using Xba/SacI and subcloned into Xba/SacI-digested pMSS vector (36), a suicide plasmid for *M. smegmatis* that contains streptomycin selection and sucrose contra-selection markers. The resultant plasmid, pMSS:ΔMSMEG_5126, was sequenced then electroporated into electrocompetent *M. smegmatis* mc^2^155 cells, using an ECM 630 electroporator (BTX), selecting for streptomycin-resistant colonies (30 μg/ml), which were then screened for sensitivity to 10% (w/v) sucrose. DNA from confirmed streptomycin-resistant, sucrose-sensitive colonies was PCR amplified using primer pair MSMEG_5126screen-F and MSMEG_5126KOright-R. The resultant PCR product was confirmed by DNA sequencing.

A confirmed single crossover (SCO) strain was grown for three days in the absence of antibiotic selection, serially diluted and plated on LB plates containing 10% (w/v) sucrose to select for potential double crossover (DCO) strains (*ie*. MSMEG_5126 deletion mutants). Genomic DNA was extracted from sucrose-resistant, streptomycin-sensitive clones, digested using ClaI/NcoI and subjected to Southern blot analysis using an MSMEG_5126 specific probe to confirm deletion of the MSMEG_5126 gene. For Southern blotting analysis 2 μg of gDNA was digested with appropriate restriction enzymes (NEB) at 37 °C for 16 hours. Purified samples and digoxygenin (DIG)-labeled, HindIII-digested λ DNA markers were separated on a 1% agarose gel followed by depurination, denaturation, neutralization, and capillary transfer onto a nylon membrane (Thermo Fisher). The membrane was then hybridized at 67 °C with a gene-specific probe prepared by DIG labelling a 3.0-kb PCR product obtained using primers MSMEG_5126screen-R and MSMEG_5126screen-F. Once confirmed by Southern blotting, the genome of a mutant was sequenced at The Peter Doherty Institute for Infection and Immunity at the University of Melbourne.

MSMEG_1829 (*fbiB*), MSMEG_2392 (*fbiD*) and MSMEG_1830 (*fbiA*) deletion mutants were generated following the same methods as MSMEG_5126, using gene-specific primer combinations (Table S3). Individual deletion mutants of *fbiA*, *fbiB* and *fbiD* were confirmed by Southern blot following digestion with PvuII and then genome sequencing.

### Purification of Fo, DH-F_420_-0 and F_420_

Fo was purified from culture supernatants of *M. smegmatis* mc^2^155 overexpressing FbiC from *M. tuberculosis* cloned into the acetamide inducible vector pMyNT. Cells were grown at 37 °C in 7H9 media to an OD_600_ ~3.0 before FbiC expression was induced by the addition of 0.2 % acetamide. Cells were grown for an additional 72 hours at 37 °C with shaking and supernatant was clarified by centrifugation at 10,000 × g for 20 minutes. The clarified supernatant was filtered (0.45 μM) and applied to a C18-silica column equilibrated in dH_2_O. Bound Fo was eluted with 20 % methanol in dH_2_O and the solvent was removed by vacuum evaporation. Fo was resuspended in dH_2_O and centrifuged (20,000 g for 20 minutes) to remove insoluble contaminants, before reapplication to a C18-silica column equilibrated in dH_2_O. Fo was again eluted with 20 % methanol in dH_2_O, vacuum evaporated and stored at −20 °C for further analysis.

F_420_ was expressed and purified as previously described in *M. smegmatis* mc24517 overexpressing FbiA, FbiB, and FbiC in the expression vector pYUBDuet-FbiABC (37). Cells were grown in LB broth + 0.05% Tween 80 at 37 °C with shaking to an OD_600_ of ~2.0 before the expression of the *fbi* genes was induced with 0.2 % acetamide. Cells were grown for an additional 72 hours before harvesting by centrifugation at 10,000 × *g* for 20 minutes. Cells were resuspended in 50 mM Tris (pH 7.5) at a ratio of 10 ml of buffer per 1 gram of wet cells and lysed by autoclaving. The autoclaved cell suspension was clarified by centrifugation at 20,000 × *g* for 20 minutes. The clarified supernatant was applied to a High Q Anion Exchange Column (Biorad) equilibrated in 50 mM Tris (pH 7.5). Bound species were eluted with a gradient of 0-100 % of 50 mM Tris, 1 M NaCl (pH 7.5). Fractions containing F_420_ were identified via visible spectroscopy based on their distinctive absorbance peak at 420 nm. Fractions containing F_420_ were pooled and applied to a C18-silica column equilibrated in dH_2_O. F_420_ was eluted with 20 % methanol in H_2_O, vacuum evaporated and stored at −20 °C for further analysis.

DH-F_420_-0 was extracted from purified FbiA expressed in *M. smegmatis* as described below. Purified concentrated FbiA (~20 mg ml^−1^; prior to cleavage of the poly-histidine tag) was denatured in a buffer containing 50 mM Tris and 8 M Urea (pH 7.0). This solution containing denatured FbiA and free DH-F_420_-0 was applied to a nickel-agarose column, with denatured FbiA binding to the column due to its hexahis-tag and DH-F_420_-0 eluting in the flowthrough. The flowthrough containing DH-F_420_-0 was applied to a Superdex 30 10/300 column equilibrated in dH_2_O and eluted fractions containing DH-F_420_-0 were identified based on their absorbance at 420 nM. DH-F_420_-0 containing fractions were pooled, vacuum evaporated, resuspended in 500 μl of dH_2_O and reapplied to the Superdex 30 10/300 column equilibrated in 20 % acetonitrile in dH_2_O. DH-F_420_-0 containing fractions were then pooled, vacuum evaporated and stored at - 20 °C for further analysis.

### FbiA expression and purification

The DNA coding sequence corresponding to FbiA from *M. smegmatis* was amplified by PCR using primers outlined in Table S3, resulting in a DNA fragment with 5’ NcoI and 3’ HindIII sites respectively. This fragment was cloned into pMyNT by restriction enzyme cloning using the aforementioned sites, yielding pMyNTFbiAMS, which expresses FbiA with a TEV cleavable N-terminal hexahistidine tag. This vector was cloned and propagated in *E. coli* DH5α in LB media/agar with the addition of 200 μg ml^−1^ hygromycin B. Sequence confirmed pMyNTFbiAMS was transformed into *M. smegmatis* mc^2^155 via electroporation, with successful transformants selected for in LB + 0.05 % Tween 80 (LBT) agar in the presence of 50 μg ml^−1^ hygromycin B. Colonies from this transformation were used to inoculate 50 ml of LBT media +50 μg ml^−1^ hygromycin B, which was grown with shaking at 37 °C until stationary phase (2-3 days). This starter culture was used to inoculate 5 liters of Terrific Broth + 0.05 % Tween 80 (TBT) giving a 1:100 dilution of the starter culture. Cells were grown with shaking at 37 °C for 24 hours until approximately mid-log phase and protein production was induced through the addition of 0.2% acetamide. Cells were grown with shaking at 37 °C for an additional 72 hours before they were harvested via centrifugation at 5,000 × *g* for 20 minutes. Harvested cells were either lysed immediately or stored frozen at −20 °C.

Cells were resuspended in Ni binding buffer (20 mM HEPES, 300 mM NaCl, 20 mM imidazole; pH 7.5) at a ratio of approximately 5 ml of buffer per 1 g of wet cell mass. Lysozyme (1 mg ml^−1^), DNAse (0.5 mg ml^−1^) and complete protease inhibitor tablets (Roche) were added and cells were lysed with a cell disruptor (Constant Systems). The cell lysate was stored on ice and clarified by centrifugation at 4 °C at 30,000 × *g*. Clarified lysate was passed through a column containing Ni^2+^ agarose resin equilibrated in Ni binding buffer. The column was washed with Ni binding buffer and protein was eluted with a gradient of Ni Gradient Buffer (20 mM HEPES, 300 mM NaCl, 500 mM imidazole; pH 7.5). Fractions containing FbiA were identified based on absorbance at 280 nm and their yellow color due to F_420_ precursor co-purification and pooled. Pooled fractions were applied a Superedex S200 26/600 size exclusion chromatography (SEC) column, equilibrated with SEC buffer (20 mM HEPES, 150 mM NaCl; pH 7.5), and fractions containing FbiA were identified as above and pooled. The hexahis-tag was cleaved from purified FbiA through the addition of 0.5 mg of hexahistidine-tagged TEV protease (expressed and purified as described in reference (38)) per mg of FbiA, plus 1 mM DTT. Digestion was performed at room temperature for ~6 hours before the sample was passed through a Ni^2+^ agarose column to remove TEV and the cleaved hexahis tag. The resulting flowthrough from this column was collected, concentrated to ~15 mg ml^−1^ and snap frozen at −80 °C. Purified FbiA was lightly yellow in color due to co-purification with F_420_ precursors, with a yield of 5-10 mg per litre of culture. The molecular weight of purified FbiA was determined by size exclusion coupled multiangle laser light scattering (SEC-MALS), using a Superdex S200 Increase 10/300 column equilibrated in 200 mM NaCl, 50 mM Tris [pH 7.9], coupled to FPLC (Shimadzu) with MALS detection (Wyatt Technology).

### FbiA crystallisation, ligand soaking and structure solution

Purified FbiA was screened for crystallisation conditions using a sparse matrix approach, with approximately 600 individual conditions screened. Thin inter-grown plate crystals of FbiA formed in a number of conditions, with a condition containing 0.1 M Tris (pH 8.0), 0.2 M Ca Acetate and 20 % PEG 3350 chosen for optimization. Diffraction quality crystals were obtained by microseeding into 0.1 M Tris (pH 8.5), 0.2 M Ca Acetate and 16 % PEG 3350 +/− 20 % glycerol. Crystals grew as bunches of very thin plates and were slightly yellow in color. Crystals from conditions containing glycerol were looped and directly flash cooled to 100 K in liquid N2, providing ‘apo’ crystals for data collection. Crystals from non-glycerol containing wells were transferred into well solution with 20 % glycerol and either Fo, GDP, Fo and GDP or DH-F_420_-0. Crystals were incubated in this solution for 1-5 minutes before they were looped and flash cooled to 100 K in liquid N_2_.

Data were collected at the Australian Synchrotron, with crystals diffracting anisotropically to ~2.2 to 3.0 Å, and processed using XDS and merged using Aimless from the CCP4 package (39, 40). The structure of FbiA was solved by molecular replacement using Phaser (41), with a search model derived from the structure of CofD from *M. mazei* (PDB ID: 3C3D) prepared based on the amino acid sequence for FbiA from *M. smegmatis* using sculptor from the Phenix package (41). Native and ligand soaked structures of FbiA were built and refined using Coot and phenix.refine from the Phenix package (41, 42). Structural coordinates for Fo, DH-F_420_-0 and F_420_ were generated using the AceDrg program within the CCP4 suite (40, 43).

### LC-MS detection of F_420_ and precursors

Wild-type and Fbi mutant *M. smegmatis* mc^2^155 strains were grown in 20 ml of LBT media until stationary phase (2-3 days) and harvested by centrifugation at 5,000 × *g* for 20 minutes. Cells were resuspended in 2 ml of dH_2_O and lysed by boiling for 5 minutes before clarification by centrifugation at 25,000 × *g* for 10 minutes. The soluble fraction was then decanted for mass spectrometry analysis. Samples were analyzed by hydrophilic interaction liquid chromatography (HILIC) coupled to high-resolution mass spectrometry (LC−MS) according to a previously published method (44). In brief, the chromatography utilized a 20 × 2.1 mm guard in series with a 150 × 4.6 mm analytical column (both ZIC-pHILIC, Merck). Column temperature was maintained at 25 °C with a gradient elution of 20 mM ammonium carbonate (A) and acetonitrile (B) (linear gradient time-%B as follows: 0 min-80%, 15 min-50%, 18 min-5%, 21 min-5%, 24 min-80%, 32 min-80%) on a Dionex RSLC3000 UHPLC (Thermo). The flow rate was maintained at 300 μl min^−1^. Samples were kept at 4 °C in the autosampler and 10 μl was injected for analysis. The mass spectrometric acquisition was performed at 35,000 resolution on a Q-Exactive Orbitrap MS (Thermo) operating in rapid switching positive (4 kV) and negative (−3.5 kV) mode electrospray ionization (capillary temperature 300 °C; sheath gas 50; Aux gas 20; sweep gas 2; probe temp 120 °C). The resulting LC-MS data were processed by integrating the area below the extracted ion chromatographic peaks using TraceFinder 4.1 (Thermo Scientific). All species were detected in negative mode as the singly deprotonated anion (Fo and DH-F_420_-0) or in the case of the F_420_-n species the double deprotonated dianion.

### 2PL synthesis

2PL was chemically synthesized using the following protocol: Benzyl lactate was condensed with chlorodiphenyl phosphate in pyridine, with cooling, to give benzyldiphenylphosphoryl lactate. Hydrogenolysis of this material in 70% aqueous tetrahydrofuran over 10% Palladium on carbon (Pd-C) gave phospholactic acid as a colorless, viscous oil, which was characterized by proton, carbon and phosphorus NMR spectroscopy, and by mass spectrometry. ^1^H NMR (DMSO-d6) δ 11.68 (br, 3H), 4.53 (m, 1H), 1.36 (d, J=6.8 Hz, 3H). ^13^C NMR (DMSO-d6) δ 172.50 (d, JP-C=0.05 Hz), 69.56 (d, JP-C=0.04 Hz), 19.27 (d, JP-C=0.04 Hz). ^31^P NMR (DMSO-d6) δ −1.64. APCI-MS found: [M+H]+=171.1, [M-H]-=169.1.

### Stimulation of DH-F_420_-0 production in spiked *M. smegmatis* cell lysates and detection of F_420_ species by HPLC

To detect F_420_ synthesis in spiked *M. smegmatis* lysates, 500 ml cultures were grown in LBT for 3 days at 37 °C with shaking. Cells were harvested by centrifugation at 8,000 × *g* for 20 min, 4 °C. The pellet was resuspended in 50 ml lysis buffer (50 mM MOPS, 1 mM phenylmethylsulfonyl fluoride (PMSF), 1 mM DTT, 5 mM MgCl^2^, 2.5 mg.ml^−1^ lysozyme, 2.5 mg Deoxyribonuclease I). An M-110P Microfluidizer (Fluigent) pressure-lysis maintained at 4 °C was used to lyse the cells. The lysate was centrifuged at 10,000 × *g* at 4 °C for 20 min. 1 ml aliquots of lysate were spiked with either 1 mM phosphate buffer (pH 7.0) plus GTP and 2PL, or GTP and PEP. These spiked samples, along with a ‘no spike’ control, were incubated at 4 hours at 37 °C. To terminate the reaction, the aliquots were heated at 95 °C for 20 min, then centrifuged at 16,000 × *g* for 10 min. The supernatants were filtered through a 0.22 μm PVDF filter and moved to analytical vials.

F_420_ biosynthetic intermediates present in the filtered *M. smegmatis* cell lysates were analysed by separation and detection using an Agilent 1200 series HPLC system equipped with a fluorescence detector and a Poroshell 120 EC-C18 2.1 x 50 mm 2.7 μm column. The system was run at a flow rate of 0.3 ml min^−1^ and the samples were excited at 420 nm and emission was detected at 480 nm. A gradient of two buffers were used: Buffer A, containing 20 mM ammonium phosphate, 10 mM tetrabutylammonium phosphate, pH 7.0. Buffer B, 100% acetonitrile. A gradient was run from 25% to 40% buffer B as follows: 0-1 min 25%, 1-10 min 25%-35%, 10-13 min 35%, 13-16 min 35-40%, 16-19 min 40%-25%.

## Supporting information

Supplemental Datafiles

## Acknowledgements

This research was undertaken in part using the MX2 beamline at the Australian Synchrotron, part of ANSTO, and made use of the Australian Cancer Research Foundation (ACRF) detector. We would like to thank the Monash Crystallisation Facility for their assistance with sample characterization, crystallographic screening, and optimization. pMyNT was a gift from Dr. Katherine Beckham, Dr. Annabel Parret, and Prof. Matthias Wilmanns at EMBL Hamburg. This work was supported by an ARC DECRA Fellowship (DE170100310; awarded to C.G.), an NHMRC EL2 Fellowship (APP1178715; awarded to C.G.), an NHMRC New Investigator Grant (APP5191146; awarded to C.G.), an NHMRC grant (APP1139832; awarded to C.J.J., C.G., and G.M.C.), a Monash University Science-Medicine Seed Grant (awarded to C.G. and M.J.C.), and Monash University Doctoral Scholarships (awarded to P.R.F.C. and D.G), an ARC Discovery Grant (DP180102463; awarded to R.L.C.) We would like to thank Dr. Ghader Bashiri for helpful discussions, Dr. Matthew Belousoff for his assistance in the development of F_420_ derivative purification protocols, and Mr. Phillip Holt in the School of Chemistry, Monash University for his assistance with Fluorescence HPLC analysis.

## Author contributions

C.G. and R.G. conceived and supervised the study. Different authors contributed to cellular spiking assays (B.N., C.G., C.J.J., A.W., M.C.T.), knockout construction (C.G., R.B., R.L.C., P.K.C., G.M.C., J.G.O., M.C.T., L.K.H., G.T., D.A.W.), LC-MS analysis (C.K.B., R.G., R.S., C.G.), protein purification (R.G., C.G., B.N., P.R.F.C., C.J.J.), and crystallographic analysis (R.G., P.R.F.C., D.L.G., C.G., T.I., M.J.C.). R.G. and C.G. analysed data and wrote the paper with input from all authors.

## References

1. Greening C, et al. (2016) Physiology, biochemistry, and applications of F420-and Fo-dependent redox reactions. Microbiol. Mol. Biol. Rev. 80(2):451–493.

2. Eirich LD, Vogels GD, & Wolfe RS (1978) Proposed structure for coenzyme F420 from *Methanobacterium*. Biochemistry 17(22):4583–4593.

3. Jacobson F & Walsh C (1984) Properties of 7, 8-didemethyl-8-hydroxy-5-deazaflavins relevant to redox coenzyme function in methanogen metabolism. Biochemistry 23(5):979–988.

4. Edmondson DE, Barman B, & Tollin G (1972) Importance of the N-5 position in flavine coenzymes. Properties of free and protein-bound 5-deaza analogs. Biochemistry 11(7):1133–1138.

5. Taylor MC, et al. (2010) Identification and characterization of two families of F420H2-dependent reductases from Mycobacteria that catalyse aflatoxin degradation. Mol. Microbiol. 78(3):561–575.

6. Li W, Khullar A, Chou S, Sacramo A, & Gerratana B (2009) Biosynthesis of sibiromycin, a potent antitumor antibiotic. Appl. Environ. Microbiol. 75(9):2869–2878.

7. Greening C, et al. (2017) Mycobacterial F420H_2_-dependent reductases promiscuously reduce diverse compounds through a common mechanism. Frontiers in microbiology 8:1000.

8. Ney B, et al. (2017) The methanogenic redox cofactor F 420 is widely synthesized by aerobic soil bacteria. The ISME journal 11(1):125.

9. Rinke C, et al. (2013) Insights into the phylogeny and coding potential of microbial dark matter. Nature 499(7459):431.

10. Wu D, et al. (2009) A phylogeny-driven genomic encyclopaedia of Bacteria and Archaea. Nature 462(7276):1056.

11. Spang A, et al. (2012) The genome of the ammonia-oxidizing Candidatus *Nitrososphaera gargensis*: insights into metabolic versatility and environmental adaptations. Environmental microbiology 14(12):3122–3145.

12. Gurumurthy M, et al. (2013) A novel F420-dependent anti-oxidant mechanism protects *Mycobacterium tuberculosis* against oxidative stress and bactericidal agents. Mol. Microbiol. 87(4):744–755.

13. Jirapanjawat T, et al. (2016) The redox cofactor F420 protects mycobacteria from diverse antimicrobial compounds and mediates a reductive detoxification system. Appl. Environ. Microbiol. 82(23):6810–6818.

14. Purwantini E & Mukhopadhyay B (2013) Rv0132c of *Mycobacterium tuberculosis* encodes a coenzyme F420-dependent hydroxymycolic acid dehydrogenase. PloS one 8(12):e81985.

15. Cellitti SE, et al. (2012) Structure of Ddn, the deazaflavin-dependent nitroreductase from *Mycobacterium tuberculosis* involved in bioreductive activation of PA-824. Structure 20(1):101–112.

16. Mohamed A, et al. (2016) Protonation state of F420H2 in the prodrug-activating deazaflavin dependent nitroreductase (Ddn) from *Mycobacterium tuberculosis*. Mol. Biosyst. 12(4):1110.

17. Lee BM, et al. (2019) The evolution of nitroimidazole antibiotic resistance in Mycobacterium tuberculosis. bioRxiv:631127.

18. Graham DE, Xu H, & White RH (2003) Identification of the 7, 8-didemethyl-8-hydroxy-5-deazariboflavin synthase required for coenzyme F 420 biosynthesis. Arch. Microbiol. 180(6):455–464.

19. Graupner M & White RH (2001) Biosynthesis of the phosphodiester bond in coenzyme F420 in the methanoarchaea. Biochemistry 40(36):10859–10872.

20. Grochowski LL, Xu H, & White RH (2008) Identification and characterization of the 2-hospho-L-lactate guanylyltransferase involved in coenzyme F420 biosynthesis. Biochemistry 47(9):3033–3037.

21. Graupner M, Xu H, & White RH (2002) Characterization of the 2-phospho-L-lactate transferase enzyme involved in coenzyme F420 biosynthesis in *Methanococcus jannaschii*. Biochemistry 41(11):3754–3761.

22. Li H, Graupner M, Xu H, & White RH (2003) CofE catalyzes the addition of two glutamates to F420-0 in F420 coenzyme biosynthesis in *Methanococcus jannaschii*. Biochemistry 42(32):9771–9778.

23. Nocek B, et al. (2007) Structure of an amide bond forming F420: γγ-glutamyl ligase from Archaeoglobus fulgidus-a member of a new family of non-ribosomal peptide synthases. J. Mol. Biol. 372(2):456–469.

24. Bashiri G, et al. (2019) A revised biosynthetic pathway for the cofactor F 420 in prokaryotes. Nat. Commun. 10(1):1558.

25. Forouhar F, et al. (2008) Molecular insights into the biosynthesis of the F420 coenzyme. J. Biol. Chem. 283(17):11832–11840.

26. Braga D, et al. (2019) Metabolic Pathway Rerouting in *Paraburkholderia rhizoxinica* Evolved Long-Overlooked Derivatives of Coenzyme F420. ACS Chem. Biol.

27. Bair TB, Isabelle DW, & Daniels L (2001) Structures of coenzyme F420 in *Mycobacterium* species. Arch. Microbiol. 176(1-2):37–43.

28. Bashiri G, et al. (2016) Elongation of the poly-γ-glutamate tail of F420 requires both domains of the F420: γ-glutamyl ligase (FbiB) of *Mycobacterium tuberculosis*. J. Biol. Chem. 291(13):6882–6894.

29. Krissinel E & Henrick K (2007) Inference of macromolecular assemblies from crystalline state. Journal of molecular biology 372(3):774–797.

30. Grinter R, Roszak AW, Cogdell RJ, Milner JJ, & Walker D (2012) The crystal structure of the lipid II-degrading bacteriocin syringacin M suggests unexpected evolutionary relationships between colicin M-like bacteriocins. Journal of Biological Chemistry 287(46):38876–38888.

31. Knape MJ, et al. (2015) Divalent metal ions Mg^2+^ and Ca^2+^ have distinct effects on protein kinase A activity and regulation. ACS chemical biology 10(10):2303–2315.

32. Braga D, et al. (2019) Metabolic pathway rerouting in *Paraburkholderia rhizoxinica* evolved long-overlooked derivatives of coenzyme F420. bioRxiv:670455.

33. Fothergill-Gilmore LA & Michels PA (1993) Evolution of glycolysis. Progress in biophysics and molecular biology 59(2):105–235.

34. Shi K, et al. (2018) T4 DNA ligase structure reveals a prototypical ATP-dependent ligase with a unique mode of sliding clamp interaction. Nucleic acids research 46(19):10474–10488.

35. Haver HL, et al. (2015) Mutations in genes for the F420 biosynthetic pathway and a nitroreductase enzyme are the primary resistance determinants in spontaneous in vitro-selected PA-824-resistant mutants of *Mycobacterium tuberculosis*. Antimicrob. Agents Chemother. 59(9):5316–5323.

36. Cashmore TJ, et al. (2017) Identification of a membrane protein required for lipomannan maturation and lipoarabinomannan synthesis in *Corynebacterineae*. J. Biol. Chem. 292(12):4976–4986.

37. Bashiri G, Rehan AM, Greenwood DR, Dickson JM, & Baker EN (2010) Metabolic engineering of cofactor F420 production in *Mycobacterium smegmatis*. PLoS ONE 5(12):e15803.

38. Tropea JE, Cherry S, & Waugh DS (2009) Expression and purification of soluble His 6-tagged TEV protease. High throughput protein expression and purification, (Springer), pp 297–307.

39. Kabsch W (2010) XDS. Acta Crystallogr. Sect. D. 66(2):125–132.

40. Winn MD, et al. (2011) Overview of the CCP4 suite and current developments. Acta Crystallographica Section D: Biological Crystallography 67(4):235–242.

41. Adams PD, et al. (2010) PHENIX: a comprehensive Python-based system for macromolecular structure solution. Acta Crystallogr. Sect. D. 66(2):213–221.

42. Emsley P, Lohkamp B, Scott WG, & Cowtan K (2010) Features and development of Coot. Acta Crystallogr. Sect. D. 66(4):486–501.

43. Long F, et al. (2017) AceDRG: a stereochemical description generator for ligands. Acta Crystallographica Section D: Structural Biology 73(2):112–122.

44. Stoessel D, et al. (2016) Metabolomics and lipidomics reveal perturbation of sphingolipid metabolism by a novel anti-trypanosomal 3-(oxazolo [4, 5-b] pyridine-2-yl) anilide. Metabolomics 12(7):126.

