## Supplemental Datafiles for "Cellular and structural basis of synthesis of the unique intermediate dehydro-F_420_-0 in mycobacteria"

**Including**

Figures S1 to S5
Tables S1 to S3
SI References

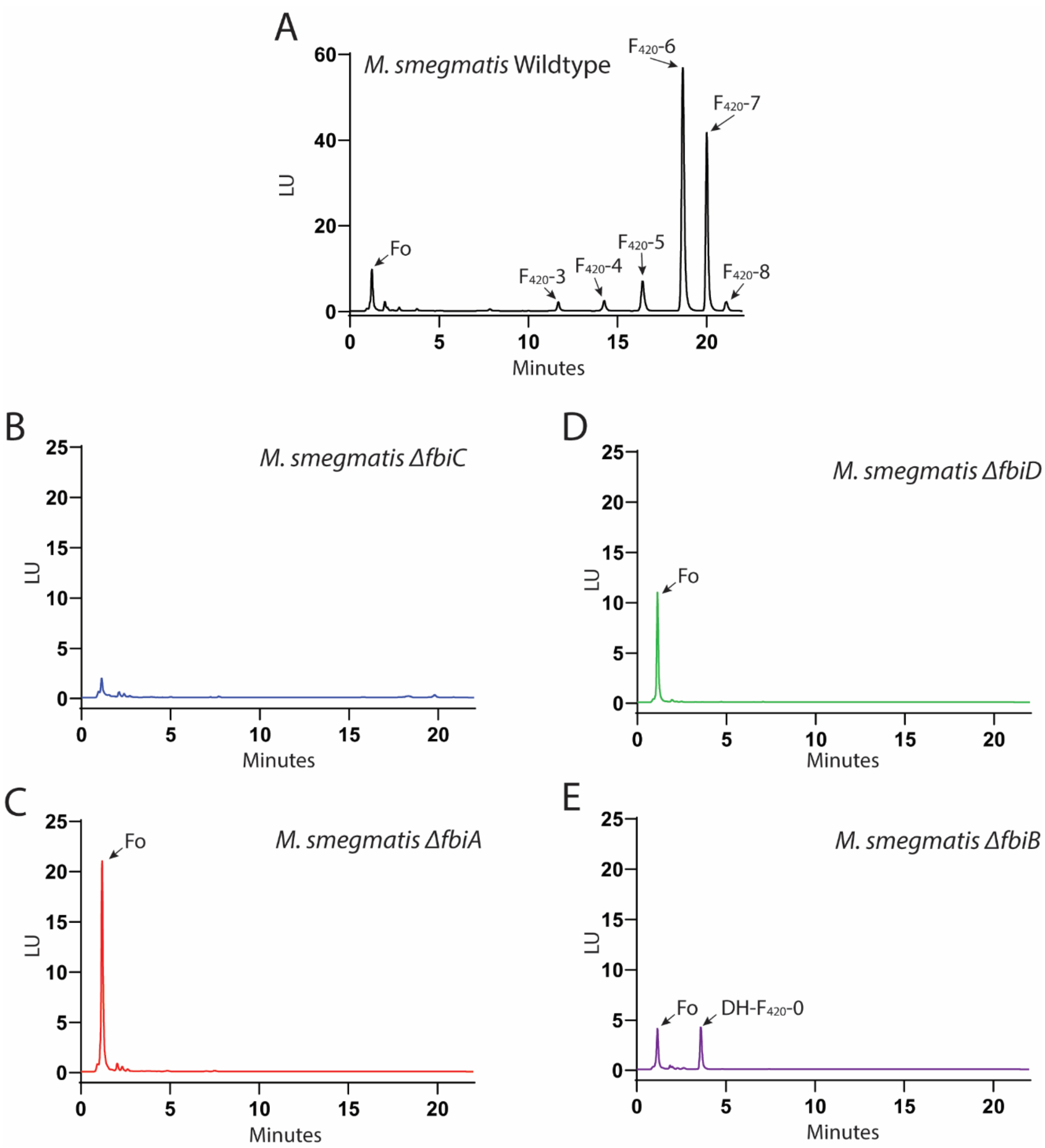


**Figure S1. Fluorescence coupled HPLC analysis of clarified cell lysates from *M. smegmatis* $F_{420}$ biosynthesis pathway mutants.** Fluorescence (Ex λ = 420 nm, Em λ = 480 nm) trace for Wildtype (A), *ΔfbiC* (B), *ΔfbiA* (C), *ΔfbiD* (D) and *ΔfbiB* (E) showing the formation of mature $F_{420}$ species in wildtype strain only and accumulation of DH-$F_{420}$-0 in *ΔfbiB* strain. LU = Fluorescence Intensity. For *ΔfbiC,* shown in panel B, mass-spectrometry analysis confirmed that no Fo or $F_{420}$ species were present, despite observed low-level fluorescence.

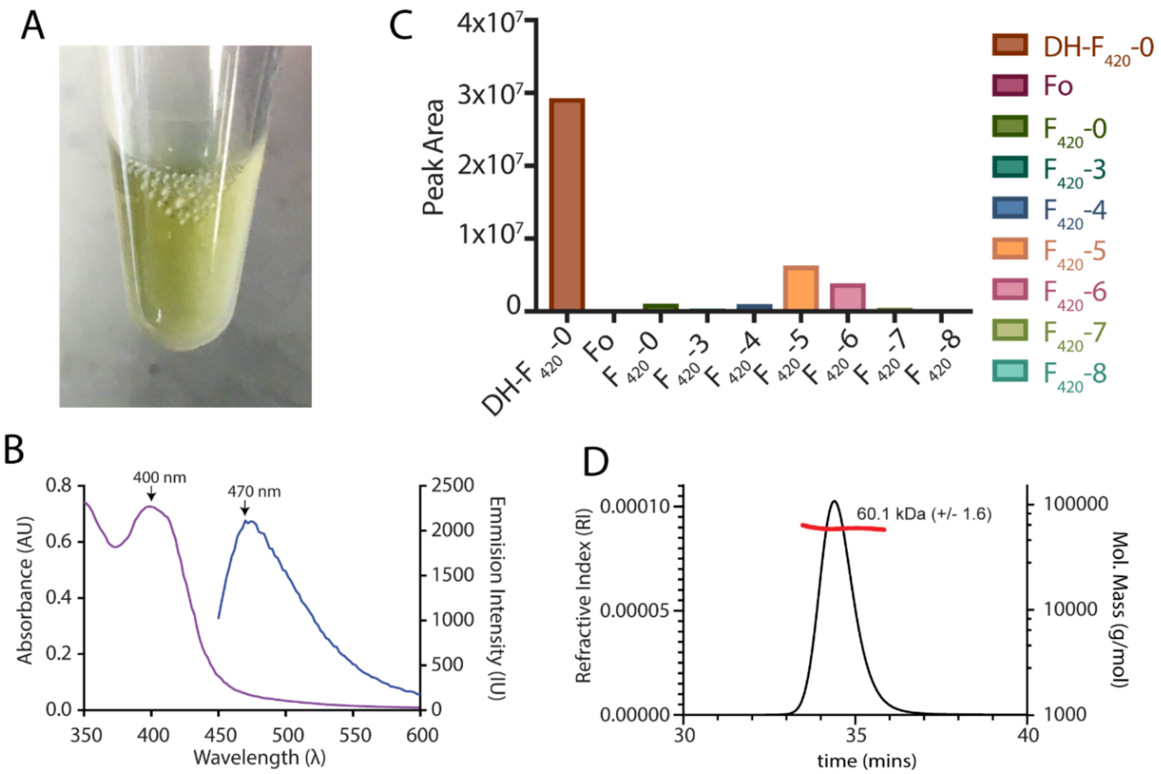


**Figure S2. FbiA recombinantly expressed in *M. smegmatis* co-purifies with its product DH-$F_{420}$-0.** (A) Purified, concentrated (16 mg.ml$^{-1}$) recombinant FbiA produced in *M. smegmatis* showing a characteristic yellow color associated with a bound $F_{420}$ species. (B) The absorbance (purple) and fluorescence spectra (blue) of purified FbiA from panel A, which is characteristic of $F_{420}$ species with a protonated deazaflavin 8-OH group. (C) LC-MS analysis of $F_{420}$ species bound to purified FbiA, showing DH-$F_{420}$-0 is the predominantly bound species and the presence of some mature $F_{420}$ species. (D) SEC-MALS analysis of purified FbiA shows that it has a molecular weight in solution consistent with a homodimeric species.

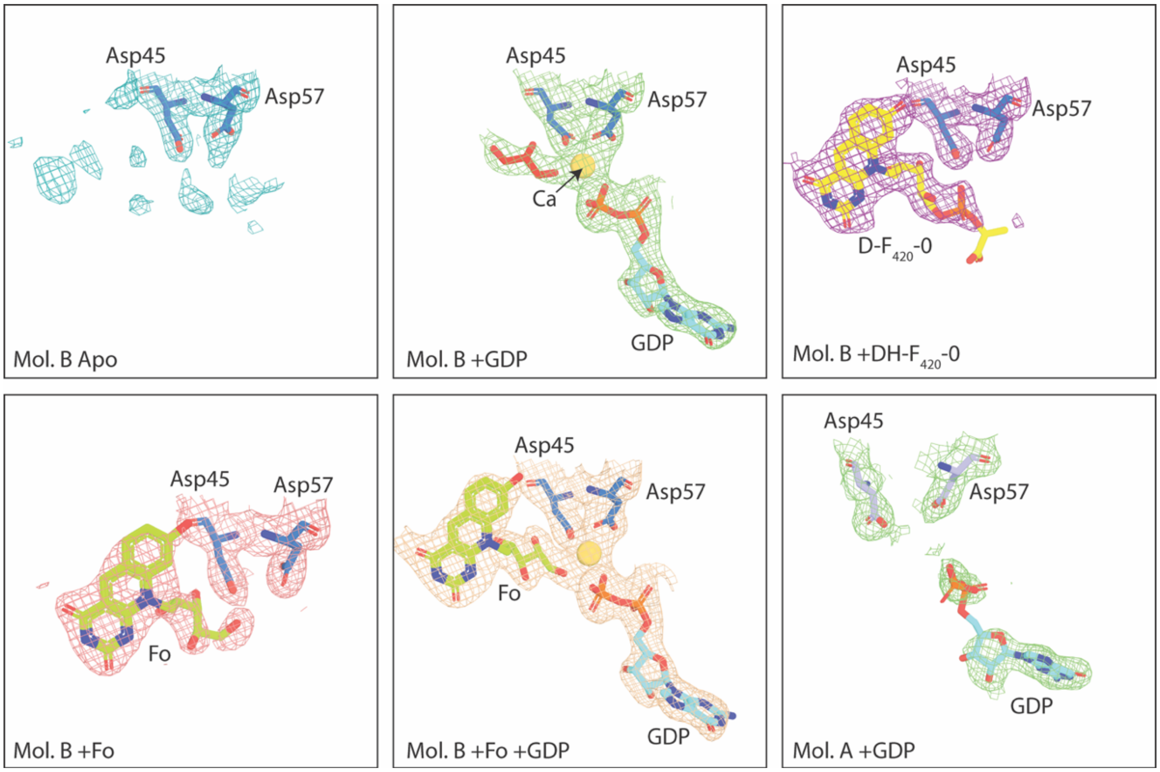


**Figure S3. Electron density corresponding to FbiA substrates and products in co-crystal structures.** Panel corresponds to the co-crystal structures indicated in the bottom right. A composite omit map is shown carved to visible molecules at a distance of 2 Å and contoured to 1 σ.

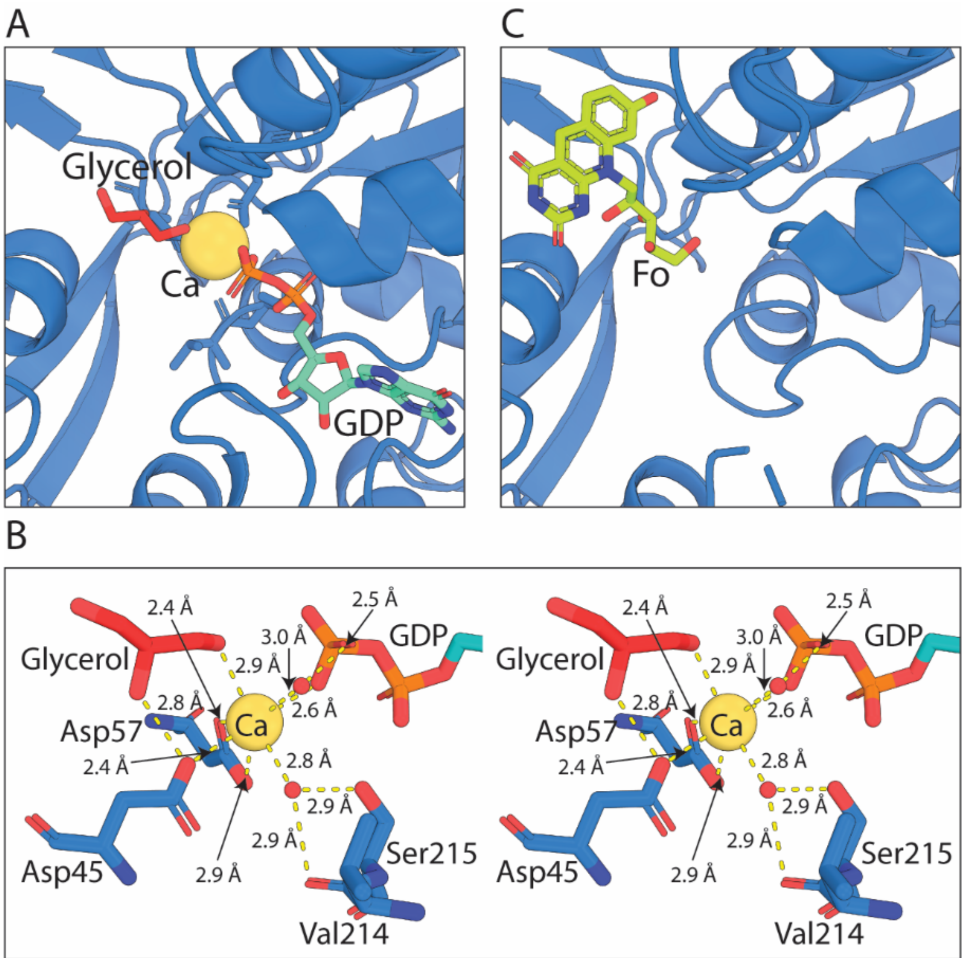


**Figure S4. The crystal structure of FbiA in complex with Fo and GDP.** (A) GDP in complex with Mol. B of FbiA. FbiA represented as a sky-blue cartoon, bound GDP and glycerol molecules shown as a stick models, bound $Ca^{2+}$ ion is shown as a yellow sphere (B) A stereoview of the catalytic center of the FbiA active site in complex with GDP and glycerol, showing FbiA sidechains involved in coordinating the catalytic metal ion and a coordinating $H_2O$ molecule. Bond distances <3.2 Å are shown as yellow dashed lines. (C) Fo in complex with Mol. B of FbiA is shown as a sky-blue cartoon, Fo as a stick model.

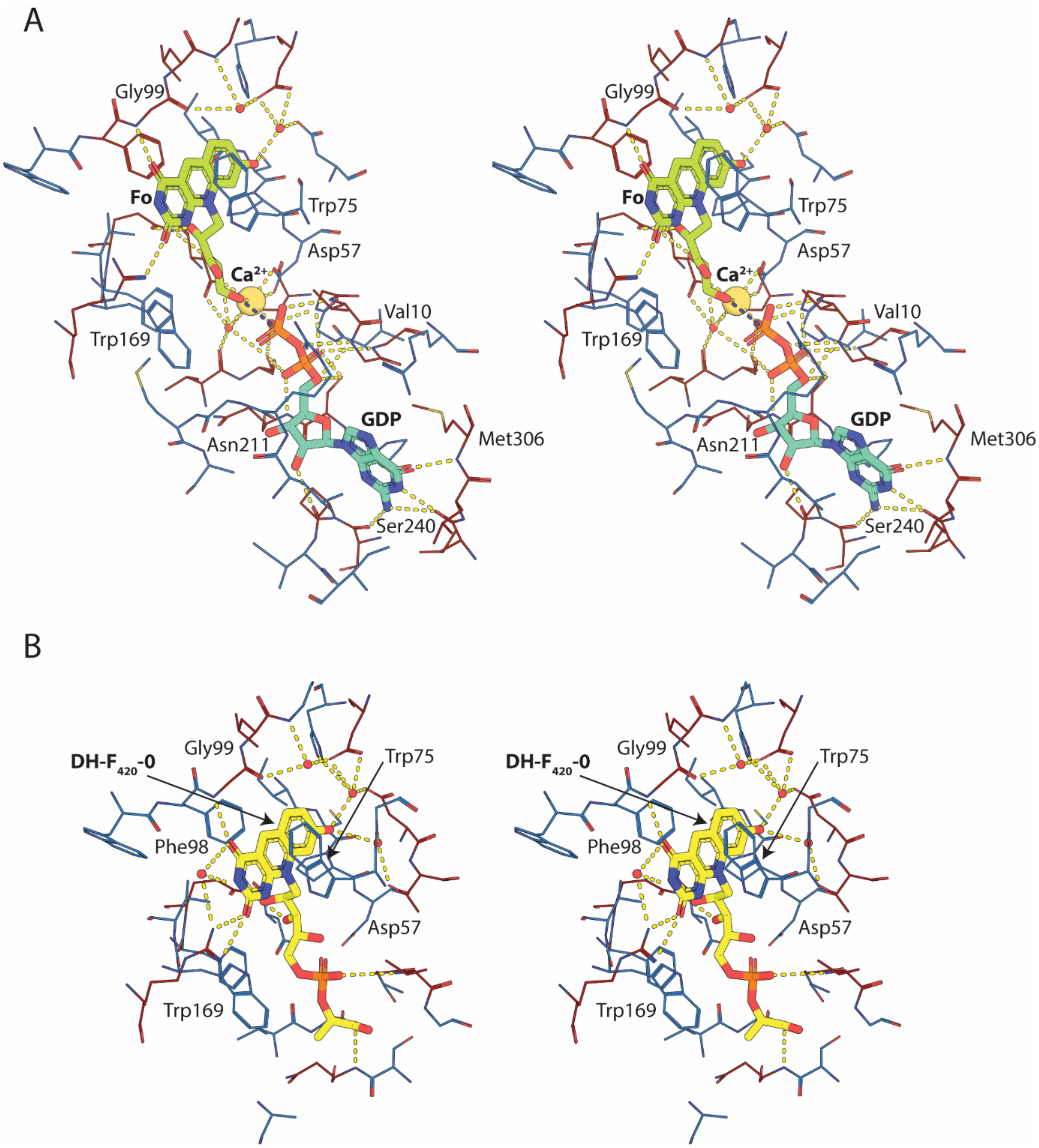


**Figure S5. The full interaction network of the FbiA active site in the presence of substrates (Fo and GDP) and product (DH-$F_{420}$-0).**

**Table S1. Crystallographic data collections and refinement statistics**

| | FbiA Apo | FbiA + Fo | FbiA + GDP | FbiA + GDP + Fo | FbiA + D-F420-0 |
|---|---|---|---|---|---|
| **Data Collection**[a] | | | | | |
| Space Group | *P2₁* → $P2_1$ | $P2_1$ | $P2_1$ | $P2_1$ | $P2_1$ |
| Cell Dimensions | | | | | |
| *a, b, c* (Å) | 46.08, 73.79, 91.47 | 46.06, 73.84, 91.70 | 46.07, 73.93, 90.93 | 46.01, 73.65, 91.27 | 46.12, 73.30, 90.34 |
| *α, β, γ* (°) | 90, 94.98, 90 | 90, 94.91, 90 | 90, 95.38, 90 | 90, 95.23, 90 | 90, 96.28, 90 |
| Wavelength | 0.976 | 0.976 | 0.976 | 0.976 | 0.976 |
| Resolution (Å) | 45.91-2.30 (2.38-2.30) | 45.90-2.20 (2.27-2.20) | 45.86-2.40 (2.49-2.40) | 45.82-2.20 (2.27-2.20) | 45.84-2.34 (2.42-2.34) |
| $R_{merge}$ | 0.215 (1.180) | 0.220 (1.612) | 0.139 (0.740) | 0.229 (2.204) | 0.124 (0.573) |
| $R_{pim}$ | 0.149 (0.841) | 0.096 (0.709) | 0.106 (0.562) | 0.168 (1.623) | 0.054 (0.252) |
| *I*/[illegible](*I*) | 6.7 (1.6) | 7.4 (1.3) | 6.7 (1.7) | 3.1 (0.7) | 12.2 (3.7) |
| *CC(1/2)* | 0.982 (0.523) | 0.990 (0.523) | 0.986 (0.644) | 0.971 (0.175) | 0.996 (0.857) |
| Completeness (%) | 99.8 (99.9) | 99.5 (94.9) | 99.6 (99.8) | 100.0 (100.0) | 99.4 (94.4) |
| Redundancy | 5.8 (5.6) | 7.1 (7.0) | 3.3 (3.2) | 3.5 (3.5) | 7.0 (7.0) |
| No. reflection | 27208 (2626) | 30908 (2545) | 23704 (2453) | 30975 (2644) | 25198 (2331) |
| **Refinement statistics** | | | | | |
| $R_{work}/R_{free}$ | 19.4/25.8 | 18.4/24.5 | 19.3/25.9 | 20.6/26.3 | 16.3/22.0 |
| No. atoms | | | | | |
| *Protein* | 4811 | 4772 | 4876 | 4823 | 4715 |
| *Ligand / ions* | 3 | 28 | 59 | 86 | 48 |
| *Solvent* | 103 | 217 | 104 | 107 | 197 |
| R.m.s deviations | | | | | |
| Bond lengths (Å) | 0.008 | 0.008 | 0.009 | 0.009 | 0.008 |
| Bond angles (°) | 1.065 | 1.048 | 1.175 | 1.168 | 1.069 |

| | | | | | |
|---|---|---|---|---|---|
| Ramachandran Plot *Favored/Allowed/Outliers (%)* | 97.95/2.05/0.00 | 97.97/1.88/0.16 | 97.99/1.85/0.15 | 97.81/2.03/0.16 | 98.23/1.77/0.00 |
| PDB ID | 6UVX | 6UW1 | 6UW3 | 6UW5 | 6UW7 |

[a] Values in parentheses are for highest-resolution shell.

**Table S2. FbiA dimer interface statistics reported by PISA[1]**

| | Mol. A | Mol. B | Inter-molecule |
|---|---|---|---|
| **Interface Atoms** | 112 | 116 | - |
| **Interface Residues** | 29 | 30 | - |
| **Buried Surface Area ($Å^2$)** | 1222.5 | 1199 | - |
| **Solvation energy ($kcal.mol^{-1}$)** | -305.4 | -321.3 | - |
| **Interface area ($Å^2$)** | - | - | 1210.8 |
| **Delta G (kcal.mol-1)** | - | - | 11.4 |
| **Binding Energy (kcal.mol-1)** | - | - | -19.3 |
| **Hydrogen Bonds** | - | - | 7 |
| **Salt Bridges** | - | - | 13 |

### Table S2. Oligonucleotide primers utilised in this study

| Primer Name | Sequence (5'-3') | Purpose |
|---|---|---|
| MSMEG_5126 left | GGCGAGCTCGCTCGCCGCGTTCTGCACCG | FbiC KO cassette generation |
| MSMEG_5126leftrev | GGCGGATCCCGGCAGCCTGCCGTTCGGAC | FbiC KO cassette generation |
| MSMEG_5126right | GGCGGATCCGTACCTGGCCGGCGGCTCCC | FbiC KO cassette generation |
| MSMEG_5126rightrev | GCTCTAGAGCTCGTTGCCCGTGTGGATC | FbiC KO cassette generation |
| MSMEG_5126screen-L | CAGCCGAGCGCGCCGTCGAC | FbiC KO cassette generation |
| MSMEG_5126KOright-R | GCTCTAGAGCTCGTTGCCCGTGTGGATC | FbiC KO cassette generation |
| MSMEG_5126screen-R | GACACCGGTGGAGACGAATG | FbiC KO cassette generation |
| MSMEG_5126screen-L | CAGCCGAGCGCGCCGTCGAC | FbiC KO cassette generation |
| MSMEG_1829KOleft-L | GGCGAGCTCGCTTGCGGCCTACGGCGTGC | FbiB KO cassette generation |
| MSMEG_1829KOleft-R | GGCGGTACCCGGAACTCGGGCAGGCCGGG | FbiB KO cassette generation |
| MSMEG_1829KOright-L | GGCGGTACCCGCGATCGGTTATCCCGAAC | FbiB KO cassette generation |
| MSMEG_1829KOright-R | GCTCTAGAGCCGGCGCGGTGATCGAACG | FbiB KO cassette generation |
| MSMEG_1829screen-L | TGCATGTACACCCTCGGCGG | FbiB KO cassette generation |
| MSMEG_1829screen-R | CCGTGGTGGGACGCGGAGCC | FbiB KO cassette generation |
| MSMEG_2392KOleft-L | GGCGAGCTCGGGTGGCGCCGAGTGCGATAC | FbiD KO cassette generation |
| MSMEG_2392KOleft-R | GGCGGTACCCGTTCTCCTGCCGCTCATGG | FbiD KO cassette generation |
| SMEG_2392KOright-L | GGCGGTACCCCCGCCACGGCACAGGCCATC | FbiD KO cassette generation |
| MSMEG_2392KOright-R | GCTCTAGATGCGACGACGATCCGAGTTC | FbiD KO cassette generation |
| MSMEG_2392screen-L | GAAGGTGCGGTTGCGGGAATG | FbiD KO cassette generation |
| MSMEG_2392screen-R | CTTCCAGTCCGGGGAACAGC | FbiD KO cassette generation |
| MSMEG_1830KOleft-L | GGCGAGCTCGATGCCGCGCTTGAGGCGCC | FbiA KO cassette generation |
| MSMEG_1830KOleft-R | GGCGGTACCCAGAACGGTGATCTTCACAAC | FbiA KO cassette generation |
| MSMEG_1830KOright-L | GGCGGTACCCGCGCCGGTCTGGACCTCGCC | FbiA KO cassette generation |
| MSMEG_1830KOright-R | GCTCTAGACGGCCTCGATGGCGTCGTGC | FbiA KO cassette generation |
| MSMEG_1830screen-L | CGTCGGTCATCGACCCGACG | FbiA KO cassette generation |
| MSMEG_1830screen-R | GGTCTCGGAGTCCTGCACCC | FbiA KO cassette generation |
| FbiA MS NcoI F | GCATCCCATGGGTAAGATCACCGTTCTGGTCGGCGG | FbiA protein expression via pMyNT |
| FbiA MS HindIII R | GATGCAAGCTTTCACAGCGACACCCCGGCGAG | FbiA protein expression via pMyNT |

**SI References**

1. Krissinel E, Henrick K. Inference of macromolecular assemblies from crystalline state. Journal of molecular biology. 2007;372(3):774-97.
